# New genes involved in Angelman syndrome-like: expanding the genetic spectrum

**DOI:** 10.1101/2020.07.16.206052

**Authors:** Cinthia Aguilera, Elisabeth Gabau, Ariadna Ramirez-Mallafré, Carme Brun-Gasca, Jana Dominguez-Carral, Veronica Delgadillo, Steve Laurie, Sophia Derdak, Natàlia Padilla, Xavier de la Cruz, Núria Capdevila, Nino Spataro, Neus Baena, Miriam Guitart, Anna Ruiz

## Abstract

Angelman syndrome (AS) is a neurogenetic disorder characterized by severe developmental delay with absence of speech, happy disposition, frequent laughter, hyperactivity, stereotypies, ataxia and seizures with specific EEG abnormalities. There is a 10-15% of patients with an AS phenotype whose genetic cause remains unknown (Angelman-like syndrome, AS-like). Whole-exome sequencing (WES) was performed on a cohort of 14 patients with clinical features of AS and no molecular diagnosis. As a result, we identified 10 *de novo* and 1 X-linked pathogenic/likely pathogenic variants in 10 neurodevelopmental genes (*SYNGAP1, VAMP2, TBL1XR1, ASXL3, SATB2, SMARCE1, SPTAN1, KCNQ3, SLC6A1* and *LAS1L*) and one deleterious *de novo* variant in a candidate gene (*HSF2*). Our results highlight the wide genetic heterogeneity in AS-like patients and expands the differential diagnosis. New AS-like genes do not interact directly with *UBE3A* gene product but are involved in synapsis and neuron system development.

## INTRODUCTION

Angelman syndrome (AS, OMIM #105830) is a neurogenetic disorder with a prevalence of about 1/15000 births. AS is characterized by severe developmental delay/intellectual disability (DD/ID) with absence of speech and distinctive dysmorphic craniofacial features such as microcephaly and wide mouth. Neurological problems include ataxia and seizures with specific EEG abnormalities. The behavioral phenotype is characterized by happy disposition, frequent laughter, hyperactivity and stereotypies [1]. The consensus criteria for the clinical diagnosis of AS was proposed in 2005 by Williams et al.,[1] which included a list of (i) consistent, (ii) frequent and (iii) associated features. However, clinical manifestations of AS can overlap with other diseases.

AS is caused by the loss of function in neuronal cells of the ubiquitin protein ligase E6-AP (E6-Associated Protein) encoded by the *UBE3A* gene, which is located on chromosome 15q11-q13 imprinted region. Methylation study of this region identifies 75–80% of AS patients including maternal deletion, paternal uniparental disomy (UPD) and imprinting center defects. Pathogenic or likely pathogenic variants in the *UBE3A* gene identify a further 10% of cases. However, for approximately 10-15% of clinically diagnosed AS patients, the genetic cause remains unknown (AS-like)[2].

Some of these AS-like patients present alternative clinical and molecular diagnoses in syndromes that have overlapping clinical phenotypes and that should be considered in the differential diagnosis of AS. AS differential diagnosis include single gene disorders such as Christianson syndrome (*SLC9A6)*, Rett syndrome (*MECP2*), Pitt Hopkins syndrome (*TCF4*), Kleefstra syndrome (*EHMT1*) and Mowat-Wilson syndrome (*ZEB2*). Individuals affected by the above mentioned syndromes present severe DD, seizures, postnatal microcephaly, absent or minimal speech and sleep disturbances as AS patients [3,4].

In order to further identify the molecular defects in AS-like patients, whole exome sequencing (WES) was performed in a cohort of 13 parent-patient trios and one single patient with clinical features of AS and no molecular diagnosis. Pathogenic/likely pathogenic variants in known neurodevelopmental genes were found in 78,5% of patients while a deleterious variant in a new candidate gene was identified in another patient. Overall, our results show that 10-15% of patients with a clinical but with no molecular diagnosis of AS present alternative genetic alterations in genes not previously associated to AS, expanding its genetic spectrum.

## MATERIALS AND METHODS

### Patient samples and clinical description of the cohort

14 patients (7 girls and 7 boys) from the Parc Taulí Hospital Universitari (Sabadell, Spain) who met the consistent clinical features of AS [1] and lacked a molecular diagnosis of AS were selected. Patient 1 had also been included in another study [5]. The corresponding informed consent was obtained from all parents for each participant in the study.

Clinical characteristics of the enrolled patients are available in Table 1. Consistent features were present in 100% of patients except for the ataxia of gait which was present in 9 of 14 patients. Even though the ataxia of gait is considered a consistent feature in AS patients a recent review shows that it ranges from 72,7% to 100 % depending on the genetic etiology [6]. All the cases were sporadic and no other relevant findings were present in their family history.

**Table 1.**
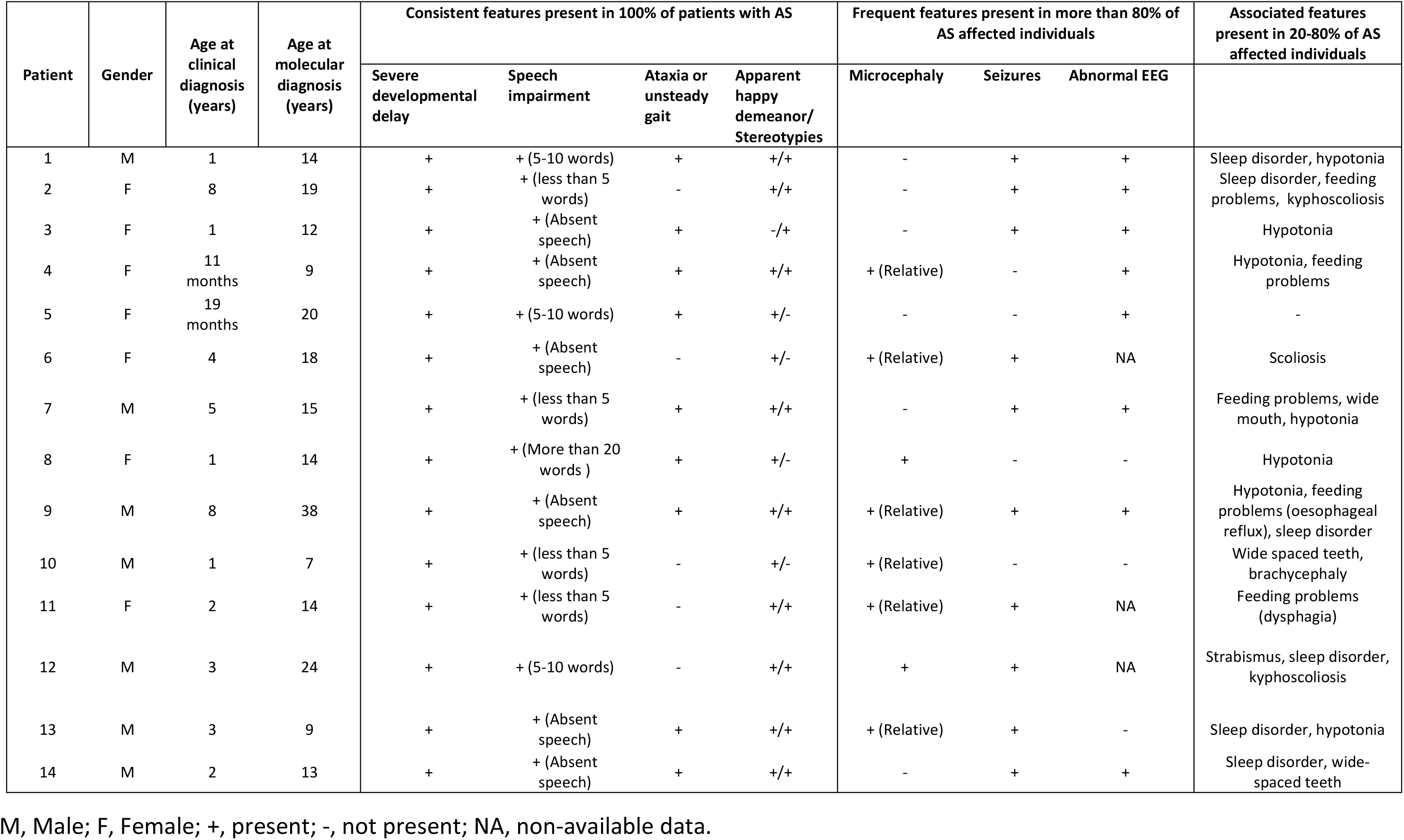
Clinical characteristics of AS-like patients.

### Whole-exome sequencing and variant interpretation

Trio WES of 13 patients and their parents was performed using the SureSelect Human All Exon V5+UTR kit (Agilent technologies). In patient 4, WES was performed only in the patient sample. Variant discovery was performed following the GATK best practices [7]. All exome variants were first checked against a *de novo* followed by an X-linked and autosomal recessive model of inheritance. The impact of missense and splice site variants was assessed using several *in silico* tools (S1 Table). Variants were classified following the ACMG guidelines [8]. Pathogenic and likely pathogenic variants have been submitted to ClinVar.

### Real time quantitative PCR (RTqPCR) analysis

RNA was extracted using the Biostic Blood Total RNA Isolation Kit sample (MO BIO laboratories, Inc) and cDNA was obtained using the PrimeScript™RT reagent Kit (Takara). RTqPCR gene expression analysis was performed using the Taqman probes HSF2-Hs00988309_g1 and GADPH-Hs02758991_g1 for normalization (Applied Biosystems).

### Network and pathway enrichment analysis in AS-like genes

Leveraging the STRING database [9], a network analysis was performed in order to examine if the new identified AS-like genes interact among themselves, with the known AS-like genes [3] or with the *UBE3A* gene. Network analysis was carried out without taking into account the text mining interaction option and using a minimum required interaction score of 0.7 (high confidence) (S1 Fig).

## Results

Identified variants were first filtered according to a dominant *de novo* model of inheritance. Variants in genes known to be involved in neurodevelopmental diseases were selected and confirmed to be *de novo*. Overall, 10 *de novo* (*SYNGAP1, VAMP2, TBL1XR1, ASXL3, SATB2, SMARCE1, SPTAN1, KCNQ3* and *SLC6A1)* and 1 X-linked (*LAS1L*) protein altering variants were confirmed in 11 patients, leading to a diagnostic yield of 78,5%. These variants were located in 10 different genes previously reported to be associated with neurodevelopmental disorders [5,10–17].

Pathogenic and likely pathogenic variants identified in this study are summarized in Table 2. Most of the genes identified in our cohort are involved in synapsis (*VAMP2, SYNGAP1, SLC6A1* and *KCNQ3*) and chromatin remodeling or transcription regulation (*TBL1XR1, SATB2, SMARCE1, ASXL3* and *LAS1L*) as has been described before in neurodevelopmental diseases [18,19].

**Table 2.** Pathogenic and likely pathogenic variants identified in AS-like patients.

| Patient | Gene | NM number | Nucleotide change | Amino acid change | Variant Type | Pattern of inheritance | Varsome Classification | Described before | Protein function |
| --- | --- | --- | --- | --- | --- | --- | --- | --- | --- |
| 1 | VAMP2 | NM_014232.2 | c.128_130delTGG | p.Val43del | In-frame | De novo | Pathogenic | Yes<br>Salpieto et al., 2019 | VAMP2 is a member of the SNARE family of proteins, which are involved in membrane fusion of synaptic vesicles. |
| 2 | SYNGAP1 | NM_006772.2 | c.1861C>T | p.Arg621* | Nonsense | De novo | Pathogenic | No | SYNGAP1 is a RAS-GTPase-activating protein with a critical role in synaptic development, structure, function and plasticity. |
| 3 | TBL1XR1 | NM_024665.5 | c.1000T>C | p.Cys334Arg | Missense | De novo | Likely pathogenic | No | TBL1XR1 is part of the repressive NCoR/SMRT complex acting as a transcriptional regulator. |
| 4 | TBL1XR1 | NM_024665.5 | c.1043A>G | p.His348Arg | Missense | De novo | Likely pathogenic | No |  |
| 5 | SATB2 | NM_001172509.1 | c.1826delA | p.Asp609Alafs*15 | Frameshift | De novo | Pathogenic | No | SATB2 participates in chromatin remodeling and transcription regulation. |
| 6 | KCNQ3 | NM_004519.3 | c.688C>T | p.Arg230Cys | Missense | De novo | Pathogenic | Yes<br>Decipher and Miceli F et al., 2015, Sands TT et al., 2019 | KCNQ3 is a voltage-gated potassium channel subunits that underlay the neuronal M-Current. |
| 7 | SMARCE1 | NG_032163.1<br>(NM_003079.4) | c.237+1G>T | p.Ala53_Lys79del | Splice site | De novo | Likely pathogenic | No | SMARCE1 is part of the SWI/SNF chromatin remodeling complex involved in transcriptional activation. |
| 8 | SPTAN1 | NM_001130438.2 | c.6592_6597dupCTGCAG | p.Leu2198_Gln2199dup | In-frame | De novo | Likely pathogenic | No | SPTAN1 is an α-II spectrin involved in stabilization and activation of membrane channels, transporters and receptors. |
| 9 | ASXL3 | NM_030632.2 | c.3106C>T | p.Arg1036* | Nonsense | De novo | Pathogenic | Yes<br>Kuechler A et al., 2016 | ASXL3 plays a role in the regulation of gene transcription and histone deubiquitination. |
| 10 | LAS1L | NM_031206.4 | c.1237G>A | p.Gly413Arg | Missense | X-linked | Likely pathogenic | No | LAS1L is involved in the 60S ribosomal subunit synthesis, maturation of 28S rRNA and regulation of transcription. |
| 14 | SLC6A1 | NM_003042.3 | c.889G>A | p.Gly297Arg | Missense | De novo | Pathogenic | Yes<br>Carvill GL et al., 2015 | SLC6A1 gene encodes for the GAT-1 GABA transporter. |

Additional clinical features of patients were analyzed taking into account the clinical phenotype described for the genes identified. The presence of specific clinical features associated to the new genes were confirmed for some of the patients. In short, cerebellar atrophy in *SPTAN1* [10], hypoplasia of the corpus callosum, hypoplasia of the 5^th^ finger nail, hypertrichosis, sparse scalp hair and aggressive behavior in *SMARCE1* [20], truncal obesity and short stature in *LAS1L* [13], myoclonic atonic seizures in *SLC6A1* [12], aggressive behavior in *SYNGAP1* [14], dysmorphic features and dental anomalies in *ASXL*3 [11] and aggressive behavior and dental anomalies in *SATB2* [16] (Fig 1).

**Fig 1.**
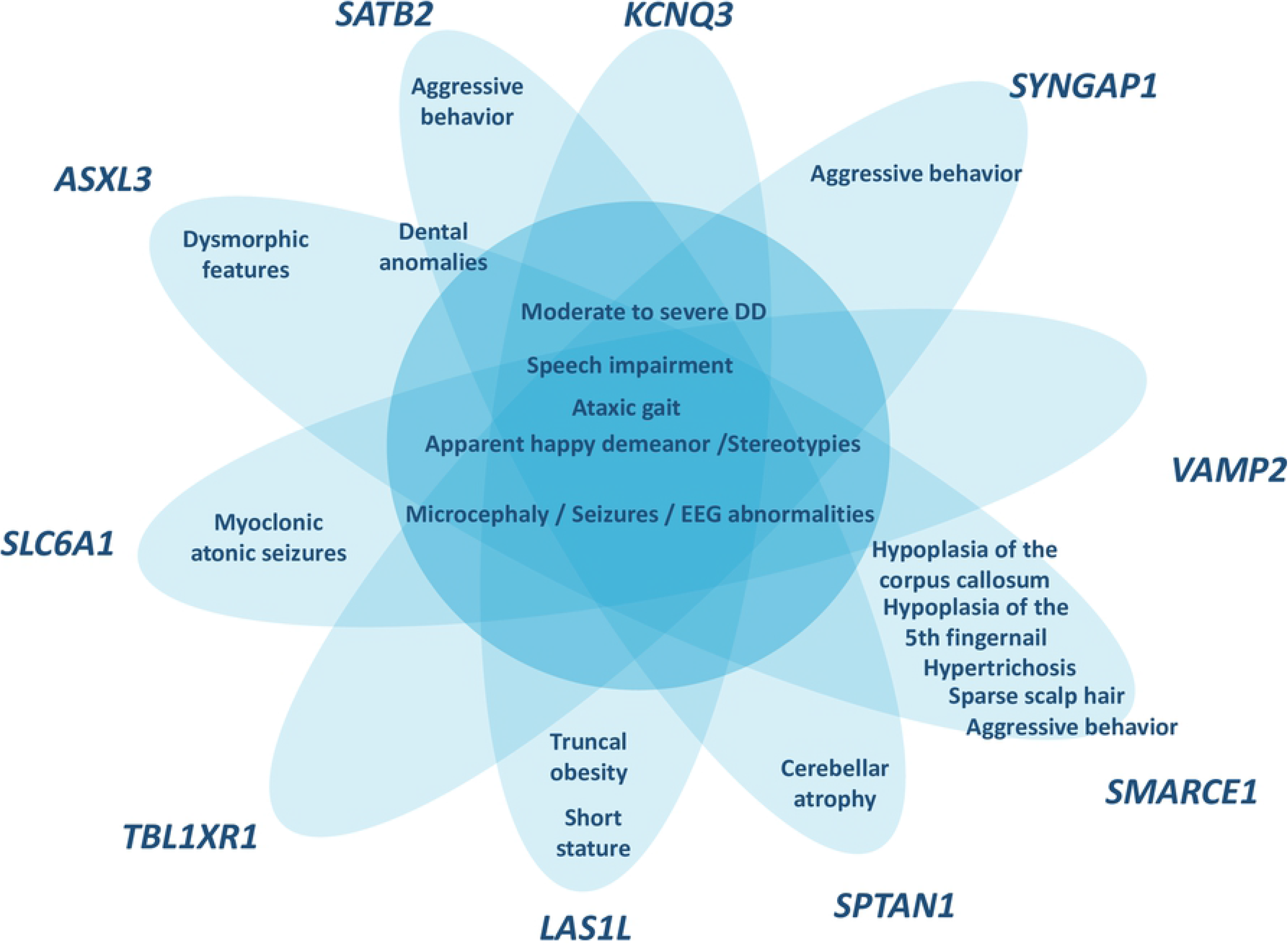
Schematic representation of the phenotypic overlap between the patients with pathogenic/likely pathogenic variants genes and the AS phenotype. In the middle of the figure there are the core features of AS that show all the patients, while in the tips there are the clinical features present in the patients of our cohort and that are associated with the gene identified.

However, not all patients presented all the clinical features associated with the genes identified. Indeed, unsteady gait and hypotonia were not present in patient carrying the pathogenic variant in *SYNGAP1* [14]; similarly the patient harboring a pathogenic variant in *SATB2* did not show sialorrhea and feeding difficulties [16]. Finally, the ataxia of gait, stereotypies and hypotonia were not observed for the patient with a pathogenic variant in *KCNQ3* [17].

A novel candidate variant was identified in a gene not previously associated with neurodevelopmental disorders. The identified variant is a *de novo* frameshift deletion c.456_459delTGAG (NM_004506.3), p.(Ser152Argfs*40) in *HSF2* gene. The variant has not been reported before and is not present in the gnomAD database. Quantification of mRNA transcripts showed a reduction in the allele carrying the frameshift variant suggesting the activation of the nonsense-mediated mRNA decay (NMD) machinery [21], supporting a loss of function mechanism of disease for the *HSF2* gene (S2 Fig).

To assess whether the genes identified in our study are functionally related to previously known AS-like genes or *UBE3A*, we tested their connectivity in the human protein-protein interaction network. No direct protein interactions were detected between the genes identified in our study, the known AS-like genes nor *UBE3A*.

However, a significant enrichment in GO terms related to the nervous system development (biological process), protein N-terminus binding (molecular function) and nuclear lumen (cellular component) were observed. Overall, these results suggest that the genes found in our cohort, the *UBE3A* gene and the known AS-like genes may not interact directly but carry out biological functions involved in synapsis and brain development, leading to overlapping phenotypes (S1 Fig).

## DISCUSSION

We identified causal variants in 11 out of 14 patients with an AS-like phenotype. The global yield diagnostic of WES in this is study is 78,5%, which is higher to what has been reported in the literature for other neurodevelopmental disorders (24-68%) [22]. The results of WES led to the identification of 10 new genes to cause an AS-like phenotype (*SYNGAP1, VAMP2, TBL1XR1, ASXL3, SATB2, SMARCE1, SPTAN1, KCNQ3, SLC6A1* and *LAS1L*), all of them previously associated with other neurodevelopmental disorders. In addition, we propose *HSF2* (Heat Shock Factor) as a new candidate gene for the AS-like phenotype. Although *HSF2* has not been previously associated with any human disease, the gene is highly expressed in the brain and highly intolerant to loss of function variation (pLI 0.92). *HSF2* knockout mice show defects in spermatogenesis and in the development of the central nervous system [23,24]. The identification of additional patients with loss of function variants in *HSF2* and functional studies in neural cells will contribute to elucidate the role of *HSF2* in the AS-like phenotype.

*De novo* variants have been described to account for approximately half of the genetic architecture of severe developmental disorders [25]. In our cohort, 10 of the 11 pathogenic and likely pathogenic variants were *de novo*, accounting for 90% of diagnosis and highlighting the power of using trio-WES for the molecular diagnosis of severe developmental disorders. Only in one case, the X-linked variant in *LAS1L* was inherited from the mother, who was a healthy mosaic carrier (20%).

All patients had received an initial diagnosis of AS, supported by the presence of consistent and frequent clinical features. In the majority of our patients (11/14) the initial diagnosis was done during infancy or early childhood (before five years old). At the time of initial diagnosis, all of them presented severe global DD and speech impairment in addition to ataxia of gait or the characteristic happy disposition. The review of patient’s clinical reports upon WES results showed the presence of additional clinical features generally not described for AS, but that were then associated with the new identified genes. Pathogenic/likely pathogenic variants in *SMARCE1, SATB2, SYNGAP1, SLC6A1, ASXL3*, *SPTAN1* and *LAS1L* genes are associated with neurodevelopment disorders that overlap with AS and with some differential features that were present in our patients (Fig 1). On the other hand, *VAMP2, KCNQ3* and *TBL1XR1* genes are associated with ID, autism spectrum disorder and epilepsy [5,15,17], features that are shared by AS-like patients.

Lack of molecular diagnosis in 10-15% of clinically diagnosed AS patients has been used to define the AS-like group. Our results indicate that the 78,5% of AS-like patients are carriers of pathogenic variants in genes involved in neurodevelopmental disorders whose features overlap with AS, showing the wide genetic heterogeneity in AS-like (Fig 1). AS-like new genes do not interact directly with *UBE3A* gene product but are involved in synapsis and nervous system development. Except for the *SYNGAP1* gene, none of the genes identified here have been previously described in the differential diagnosis of AS [26]. We propose the genes identified in this study should be included in the AS differential diagnosis and that trio WES should be considered as first line approach for the molecular diagnosis of AS-like patients. A high rate of diagnosis in patients with AS-like is essential, contributing to more appropriate clinical patient surveillance as well as allowing family genetic counseling.

## Acknowledgements

We thank the patients and families for their participation in this study.

## Supporting information

**S1 Fig. Functional and enrichment network analysis of protein-protein interactions of the candidate genes identified in our cohort, the known AS-like genes and *UBE3A*, using STRING. A)** The figure shows the interaction pattern between the different genes; circumferences and edges represent the genes and their interactions, respectively. Edges are colored according to whether there is a known interaction, a predicted interaction or others. **B)** The table shows the GO terms significantly represented amongst the candidate genes. The analysis is repeated for the three GO domains: Biological Process, Molecular Function and Cellular Component. Overall, we see that the significant terms refer to different aspects of the nervous system, from development to synapses.

**S2 Fig. Quantification of *HSF2* mRNA transcripts suggest that variant c.456_459delTGAG is sensitive to NMD. A)** qPCR analysis of *HSF2* gene expression in patient 13 and a control sample normalized to GAPDH shows less *HSF2* expression in patient 13 (* p-value 0.014). **B)** Sanger sequencing of a fragment encompassing variant c.456_459delTGAG from patient 12 and a control sample shows a reduction in the percentage of the allele with the variant in the cDNA compared to DNA.

